# Thermophilic traits correlate with slow growth in permafrost soils

**DOI:** 10.1101/2025.07.23.666387

**Authors:** Iyanu Oduwole, Tatiana A. Vishnivetskaya, Andrew D. Steen

## Abstract

Permafrost soil is characterized by prolonged freezing conditions. Thermophilic microbes have been discovered in various permanently cold environments, including permafrost, where they can persist for extended periods. The reason for this apparent mismatch between microbial adaptations and environmental conditions is unclear. Here, we test the hypothesis that thermophilic traits provide selective advantage to extremely slow-growing microbes, even in cold temperatures. We used a computational approach to predict optimal growth rates and several measures of thermophilicity in metagenome-assembled genomes (MAGs) from permafrost and active layer soils in diverse cold regions. We find that in permafrost, where available energy is always low, measures of thermophilicity correlate positively with minimum doubling time, indicating that slow growers in permafrost have more thermophilic traits. This trend is reversed in microbes in active layer soil, in which seasonal thawing, temperature changes, and episodic rain events allow periodic fast growth. Similar trends were observed in the relationship between optimal growth rates and the optimal temperature of nucleoside diphosphate kinase (NDPK), an enzyme whose temperature optimum is known to be correlated to optimal growth temperatures of the host organism. Thermophilic traits within slow growers appear to be environmentally rather than phylogenetically constrained, and thermophilic slow growers share few horizontal gene transfers with other permafrost microbes. These findings suggest that the presence of thermophilic traits in slow-growers appears to be an adaptation to extreme slow growth in a persistently low-energy environment.

**Importance:** Permanently cold environments, including permafrost soils, contain an active microbial community, which appears to include thermophilic, or heat-loving, microorganisms. This appears to be a paradox – how (and why) do microbes adapted to high temperatures live in permanently cold environments? We provide a potential answer: that the well-understood adaptations which allow microorganisms to survive high temperatures are similar to the poorly understood adaptations that allow microbes to persist over long timescales in very low-energy environments, including permafrost and the Earth’s deep subsurface. The latter environments represent 88% of the all biomass of bacteria and archaea on Earth, but the adaptations of deep subsurface microorganisms are poorly understood. This work is a step towards understanding how microorganisms persist in two different, challenging environments.

## Introduction

Permafrost soils, by definition, remain frozen for at least two consecutive years. They are often overlain by the active layer, which consists of the soils that freeze and thaw seasonally (1). Despite the inhospitable conditions, a diverse community of microbes remains active in both permafrost and active layer soils (2–4). Oddly, some of these microbes appear to be thermophilic, even though these environments do not experience high temperatures.

Several studies have reported the presence of thermophiles (organisms that grow at temperatures above 40°C) in permafrost, seasonally frozen active layer soils and other permanently cold environments such as marine sediments with temperature in range of -2° and <15°C (5–8). Many previous studies have focused on spore-forming thermophiles living in hot environments, and the ability of those microorganisms to survive in dormant state in the cold (5, 7, 9). However, recent 16S rRNA gene studies indicate the presence of non-spore-forming thermophiles in permanently cold environments (8, 10). Simultaneous incubation of permafrost samples at a variety of temperatures and in diverse media showed the presence of multiple thermophilic microorganisms. These collectively represented 1% to ∼47% of the total cultivable microbial community for both aerobic heterotrophic bacteria, which grew at 55°C, and anaerobic microorganisms, which were able to grow at 75°C (8). The latter results provide evidence that thermophiles might survive in permafrost over long timescales and become active when conditions are favorable.

Understanding how these apparent thermophiles survive in permafrost is challenging, because, in general, thermophiles do not grow in the cold (11–13). Previous studies have shown that hyperthermophiles stored in cold sea water can survive up to nine months under laboratory conditions (14), suggesting that thermophilic microbes may be transported from hot environments to cold soils. However, permafrost represents a uniquely challenging environment, including sub-freezing temperatures, low water activity, elevated salinity, and limited nutrient availability (15) which demand specific adaptations for microbes to persist (16–18). The presence of apparent thermophiles therefore seems unusual. We hypothesize that the thermophily, or thermophilic-like traits, of certain permafrost microbes represent an adaptation that facilitates an ecological strategy of extremely slow growth at consistently low temperatures.

Computational tools can successfully predict microbial physiological parameters, including optimal growth temperature (OGT) and minimum doubling time, from microbial genomes (19–21). To test whether thermophilic-like traits are an adaptation to a slow-growth life strategy, we computationally predicted traits associated with thermophiles, including optimal growth temperature, thermophilic index in genome-wide amino acid composition, and optimal temperatures of the enzyme nucleoside diphosphate kinase (NDPK), for metagenome-assembled genomes (MAGs) obtained from various permafrost and active layer environments. We also predicted organismal minimum doubling time (i.e., doubling time under optimal growth conditions in pure culture), since long minimum doubling times are a proxy for microbes with an evolutionary strategy oriented towards extreme slow growth (22). We compared these to microbes in the seasonally thawed layer of soils, so called active layer, overlying permafrost in which microbes are periodically able to metabolize and grow rapidly (23–26). Specifically, we aim to: (1) test whether there is a correlation between minimum doubling time and optimal growth temperature, and whether any correlation differs between permafrost and active layer; (2) ascertain whether phylogeny or environmental factors shape these traits; and (3) investigate the role of horizontal gene transfer in conferring traits to apparent thermophiles in permafrost soils. This study enhances our understanding of the evolutionary and adaptive strategies of permafrost microbes by investigating the thermophilic traits as a potential mechanism for sustaining slow growth in cold environment.

## Methods

### Environmental description

The permafrost samples analyzed in this study were collected from various locations with distinct thermal profiles. Siberian permafrost, a "cold" perennially frozen soil, maintains subzero temperatures between -12°C and -8°C at depths of 3-25 meters (27). Permafrost core samples were from borehole AL1_15 with total depth of 25.8 m (28), borehole Ch1_17 with total depth of 21.5 m (29), and borehole AL3_15 with total depth of 20 m (27), which were collected within the Northeastern Kolyma-Indigirka Lowland Region in Siberia, Russia between 2015-2017 and described in previous studies(27–29). The active layer samples originated from Svalbard and Stordalen Mire. By contrast, Svalbard permafrost has an average temperature of -2.8°C and is classified as "warm" permafrost (30). Svalbard active layer samples were collected at Bayelva, Ny Ålesund, Svalbard, from depths of 21–58 cm (31). Stordalen Mire, a discontinuous permafrost region with active layer soil temperatures ranging from 3.5°C to 11°C (32), soil samples were collected from 1-51 cm depth (33). Alpine permafrost and active layers were collected from the Tibetan Plateau, featuring a mean annual temperature of -4.5°C to 1.8°C; active layer samples were taken from 0-10 cm, and permafrost samples from 150-200 cm (34).

### Metagenome data sets

The permafrost datasets included 125 metagenome-assembled genomes (MAGs) from Siberia (60 from Wu et al., 52 from Liang et al., and 35 from Sipes et al.)(27–29) and 35 MAGs from alpine region of the Tibetan plateau. Active layer samples included 167 MAGs from Svalbard (27) and 507 from Stordalen Mire (33) and 24 MAGs from the Alpine region.

Additionally, we included 957 MAGs from human gut samples (35). For Alpine MAGs, 66 metagenomic sequence data from permafrost and active layer samples were downloaded from PRJNA1037019 (34). The raw metagenomic reads were quality-filtered using Trimmomatic (36), assembled with MEGAHIT (37), and binned into MAGs using MaxBin2 (38), CONCOCT (39), and MetaBAT2 (40), followed by dereplication with DAS Tool (41), checked for completeness and contamination with CheckM (42), and taxonomical classified with GTDB-TK (43). All these analyses were done on the KBase platform and are available under these narratives (https://narrative.kbase.us/narrative/195818, https://narrative.kbase.us/narrative/191544 and https://narrative.kbase.us/narrative/193190). All MAGs have >50% completeness and <10% contamination. Protein-coding open reading frames (ORFs) in the MAGs were annotated using Prokka 2.6.3 (44). We searched and filtered Nucleoside diphosphate kinase sequences from the genome and confirmed the annotation with Pfam. Genes for energy production, and conversion, and carbohydrate transport/metabolism genes were annotated using eggNOG-mapper (45).

### Growth rate Prediction

The bacterial minimum doubling time (a proxy for growth rate) was estimated with gRodon (21), an R package that estimate maximum growth rate from genomic data using codon usage bias. For the Prokka, annotated ribosomal proteins were filtered using the search “ribosomal protein”. Ribosomal proteins are often used as the set of highly expressed genes in gRodon prediction because they are known to have higher translational efficiency and universal across the tree of life (46, 47). Because our study is an extreme environment, the Madin et al. dataset (48) was selected as the training set of gRodon prediction. This phenotypic dataset included more extremophiles than that complied by Vierira-Silva and Rocha used originally in GrowthPred (49). The default minimum gene length of 240 bp was used. Despite gRodon’s tendency to underestimate slow growers’ rates due to transcription/translation efficiency biases, it has higher accuracy with the slow growers compared to other growth rate predictors (50) We expect a reliable estimation across permafrost samples since it reflects the overall microbial life style.

### Optimal growth temperature and optimal temperature of enzymes and thermophilicity prediction

Optimal growth temperature (OGT) was predicted using TOME (Temperature Optimal for Microorganisms and Enzymes) (19), which is a tool developed by training a machine learning model on amino acid composition and dipeptide frequencies of 21498 microorganisms and their OGT. TOME’s suitability for predicting thermophiles makes it ideal for this dataset. For the optimal temperature of Nucleoside Diphosphate Kinase (NDPK), the Tomer, an extended version of TOME, with improved accuracy from resampling was used (51). The thermophilic index was computed by taking the sum fraction the hydrophobic amino acid set consisting of Ile, Val, Tyr, Trp, Arg, Glu, Leu ((IVYWREL) in the amino acid sequences of each MAG (52). The same analysis was also conducted for COG functional genes related to energy production and conversion (COG C) and carbohydrate metabolism and transport (COG G) of each MAG.

### Phylogenetic signals and regression

To explore phylogenetic influences on predicted OGT and predicted minimum doubling time, a phylogenetic tree in IQTREE-2 (53) was generated using 16S rRNA sequences extracted with Barrnap (https://github.com/tseemann/barrnap). The phylogenetic signal (λ) was computed for each trait using the phylosig function in package phytools (version 0.7-70) (54) and conducted phylogenetic generalized least squares (PGLS) regression with the phylolm package (version 2.6.2) using the default parameters (55). P-values were adjusted for false positives using the Benjamini-Hochberg (BH) method with the p.adjust function in R.

### Horizontal gene transfer prediction

MAGs were classified into four categories based on the predicted OGT and minimum doubling time: thermophilic slow growers (TS; OGT > 40°C, doubling time > 5 hrs), thermophilic fast growers (TF; OGT > 40°C, doubling time < 5 hrs), mesophilic slow growers (MS; OGT 20-40°C, doubling time > 5 hrs), and mesophilic fast growers (MF; OGT 20-40°C, doubling time < 5 hrs). The 5 hrs minimum doubling time cut off for slow and fast growers was based on gRodon (21). Putative horizontal gene transfer (HGT) events were predicted using MetaCHIP (56), employing phylogenetic and best match approaches. Gene flow between categories was visualized using the R circlize package.

## Results

### Slow growers in permafrost and the active layer

We used gRodon (21) to computationally predict the maximal growth rate - i.e., the fastest growth rate possible under optimal growth conditions calculated as the minimum doubling time - for metagenome-assembled genomes (MAGs) binned from permafrost and active-layer soils from the same alpine permafrost site on Tibetan Plateau (34). This data set had a relatively low number of MAGs (35 and 24, respectively), and we were unable to identify other sites at which MAGs had been binned from both permafrost and overlying active layer soils, so we also selected two active layer sites with numerous, high quality MAGs (Ny-Ålesund, Svalbard and Stordalen Mire palsa, Sweden) plus one permafrost site with comparable quality MAGs (Kolyma-Indigirka Lowland, Siberia).

Perhaps unsurprisingly, MAGs in both permafrost and active layer soils were predicted to grow significantly more slowly than human gut MAGs (Fig. S1). There were differences in predicted growth rate among all permafrost and active layer sites, but these differences were small and unlikely to be meaningful in the real world. gRodon is not able to quantitatively predict maximal growth rates above a doubling time of ∼five hours, and in all soil samples, the predicted median growth rate was above 5.33 hours, so these results do not necessarily support actual differences in the maximal growth rates of microbes among these sites (Fig. S1, Fig. 2a).

### Thermophilicity is Positively Correlated with Doubling Time in Permafrost But Not in Active Layer Soils

Optimal growth temperature (as predicted by TOME, (19)) and “thermophilic index” (i.e., the fraction of codons in the genome coding for isoleucine, valine, tyrosine, tryptophan, arginine, glutamic acid, and leucine (52) were positively correlated to the natural log of predicted minimum doubling time in permafrost MAGs, indicating that slower growers were adapted to higher temperatures, and negatively correlated in the active layer. Only one of these correlations was statistically significant at α=0.05: the positive correlation between predicted growth temperature and minimum doubling time in alpine permafrost (p=0.0024, R=0.50, n=35) (Fig. 1a).

**Fig. 1.**
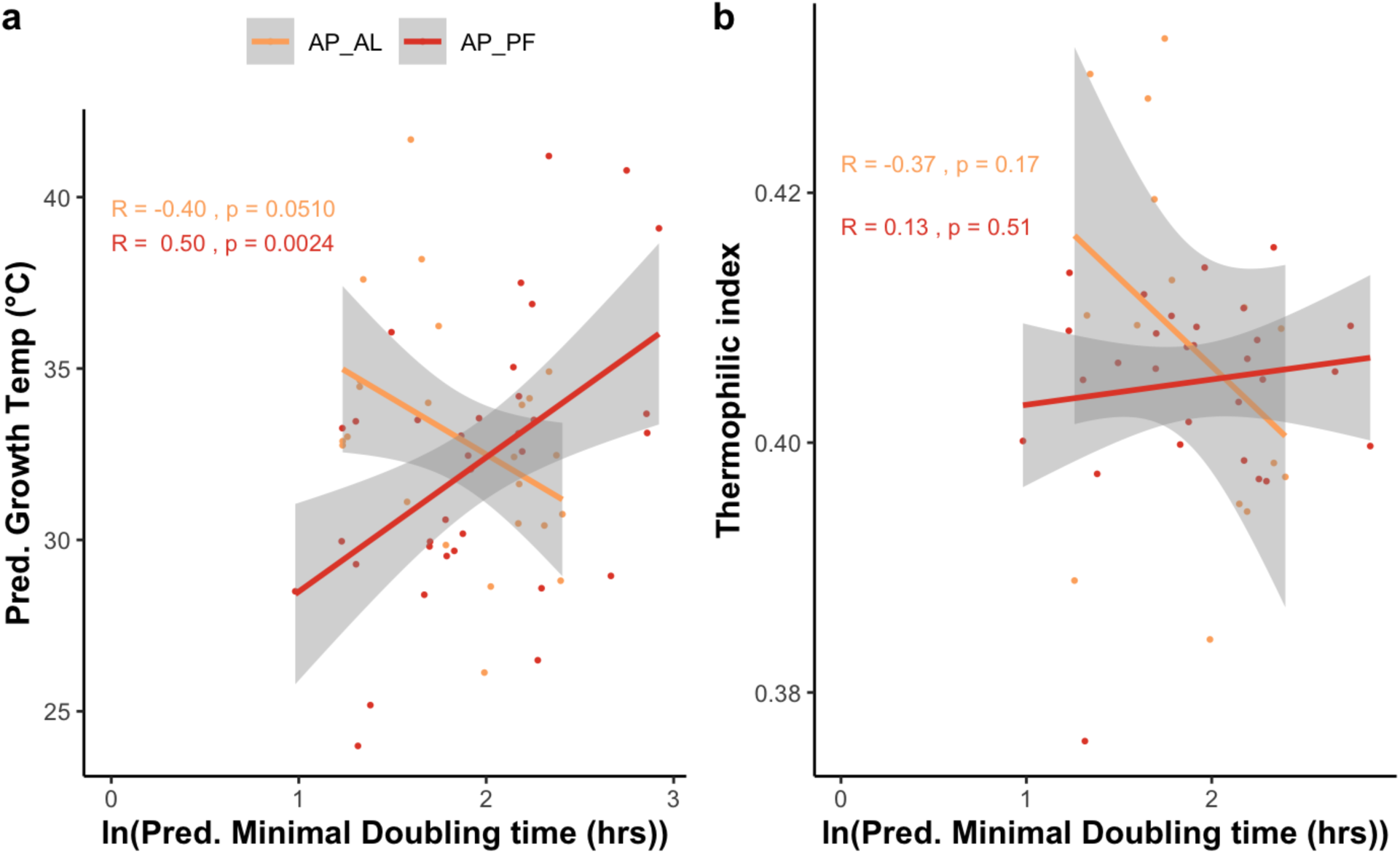
Correlations between predicted minimum doubling time by gRodon [21] and either (a) predicted growth temperature by TOME [19]; or (b) thermophilic index in permafrost and active layer from the Tibetan Plateau alpine region. Thermophilic index is the fraction amount of amino acid set: Ile, Val, Tyr, Trp, Arg, Glu, Leu (IVYWREL) which serves as indicator for thermophilicity and thermostability Abbreviations: AP_AL - alpine active layer; AP_PF - alpine permafrost.

Statistically significant correlations in the same directions were observed in all other permafrost and active layer soils (Fig. 2). In permafrost, faster-growing microbes (lower predicted minimum doubling times) tended to have lower optimal growth temperatures and thermophilic index, whereas in both active layer soils, faster growers tended to have higher predicted optimal growth temperatures and thermophilic index (regression statistics are noted in Fig. 2b). We predicted the minimum doubling times of 125 MAGs from Siberian permafrost.

**Fig. 2.**
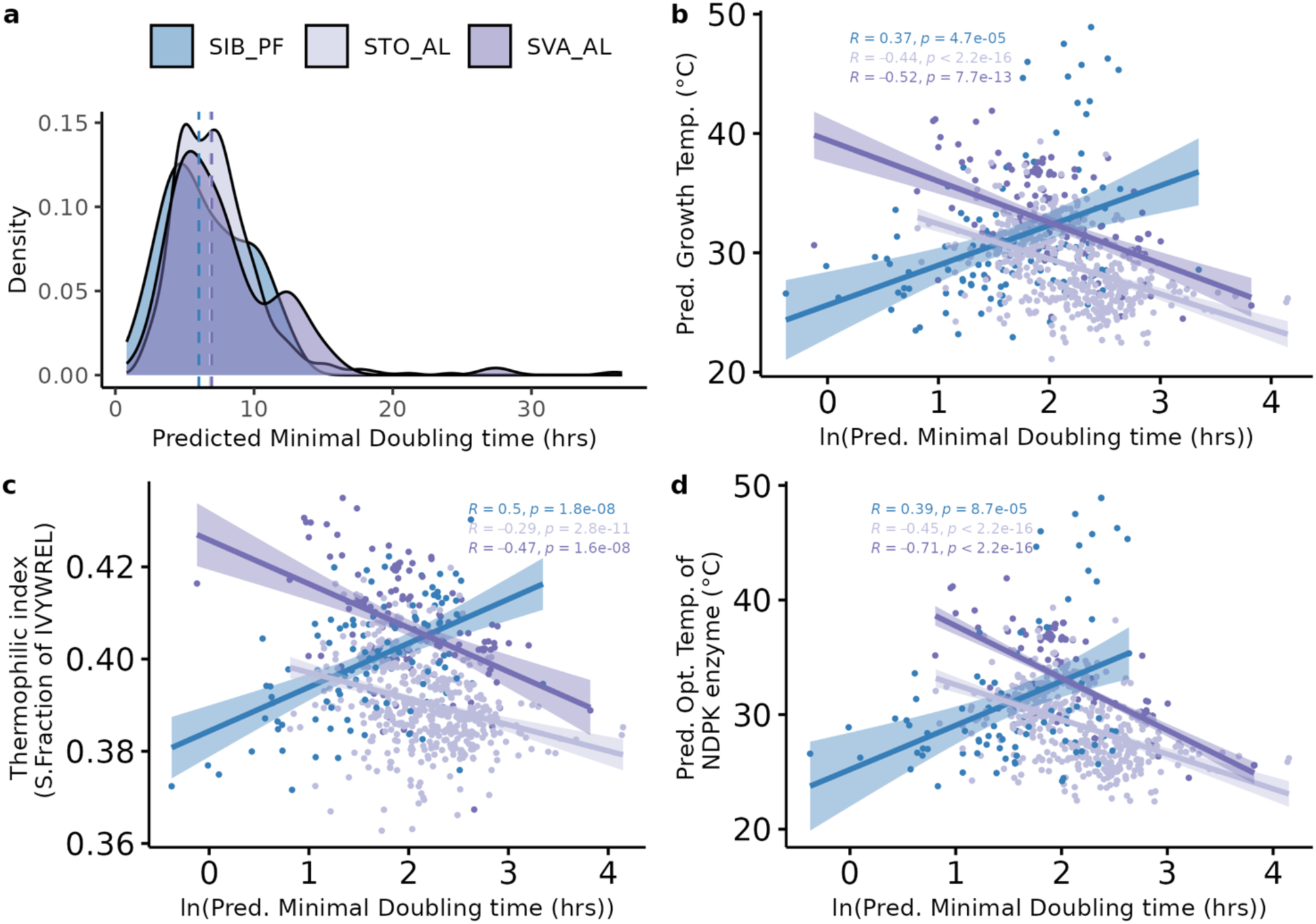
Thermophilicity is correlated with minimum doubling time. (a) Predicted minimum doubling time (hrs) of microbes from permafrost and active layer using gRodon [21]. The vertical dashed lines represent the median of the predicted minimum doubling time. Abbreviation as follows: SVA_AL - Svalbard active layer; STO_AL - Stordalen Mire palsa active layer; SIB_PF - Siberian permafrost. (b-d) Relationship between thermophilicity and growth rate in permafrost and active layer soils. Thermophilicity was measured by predicted optimal growth temperature as predicted by TOME [19] (b), by the optimal activity temperature for NDPK as predicted by TOMER [29], and as the thermophilic index. All correlation analyses were two tailed and the shaded areas around the regression line represent the 95% confidence interval.

Because data sets with substantial numbers of MAGs from corresponding permafrost soils and active layers are rare, we compared these predictions to 676 MAGs from two active layer soils: 167 MAGs from Svalbard and 507 MAGs from Stordalen Mire palsa. For these samples, we also predicted the optimum temperature of nucleoside diphosphate kinases (NDPK), a ubiquitous enzyme that catalyzes phosphate transfer from nucleoside triphosphate to nucleoside diphosphate (Fig. 1d). The same correlations were observed for all three proxies for growth temperature.

### Thermophilicity of functional genes is correlated to slow growth in permafrost

We hypothesized that functional genes, especially those related to carbon cycling, would reflect more thermophilic traits, particularly in slow-growing microbes from permafrost. Our analysis focused on genes involved in energy production and conversion (COG category ’C’) and carbohydrate transport and metabolism (COG category ’G’), as these play critical roles in soil carbon cycling. Minimum doubling times and thermophilicity were positively correlated for both gene categories (COG category C: R = 0.35, p < 0.05; COG category G: R = 0.25, p < 0.05; Fig 3), suggesting that enhanced stability of these functional genes may support microbial carbon cycling at slow growth rates. Furthermore, there was a negative correlation between minimum doubling time and the abundance of these genes (COG category C: R = -0.45, p < 0.05; COG category G: R = -0.35, p < 0.05; Fig. S2.) This indicates that slower-growing microbes have fewer genes associated with energy production and carbohydrate metabolism, yet the genes they possess code for enzymes that are more thermostable.

**Fig. 3:**
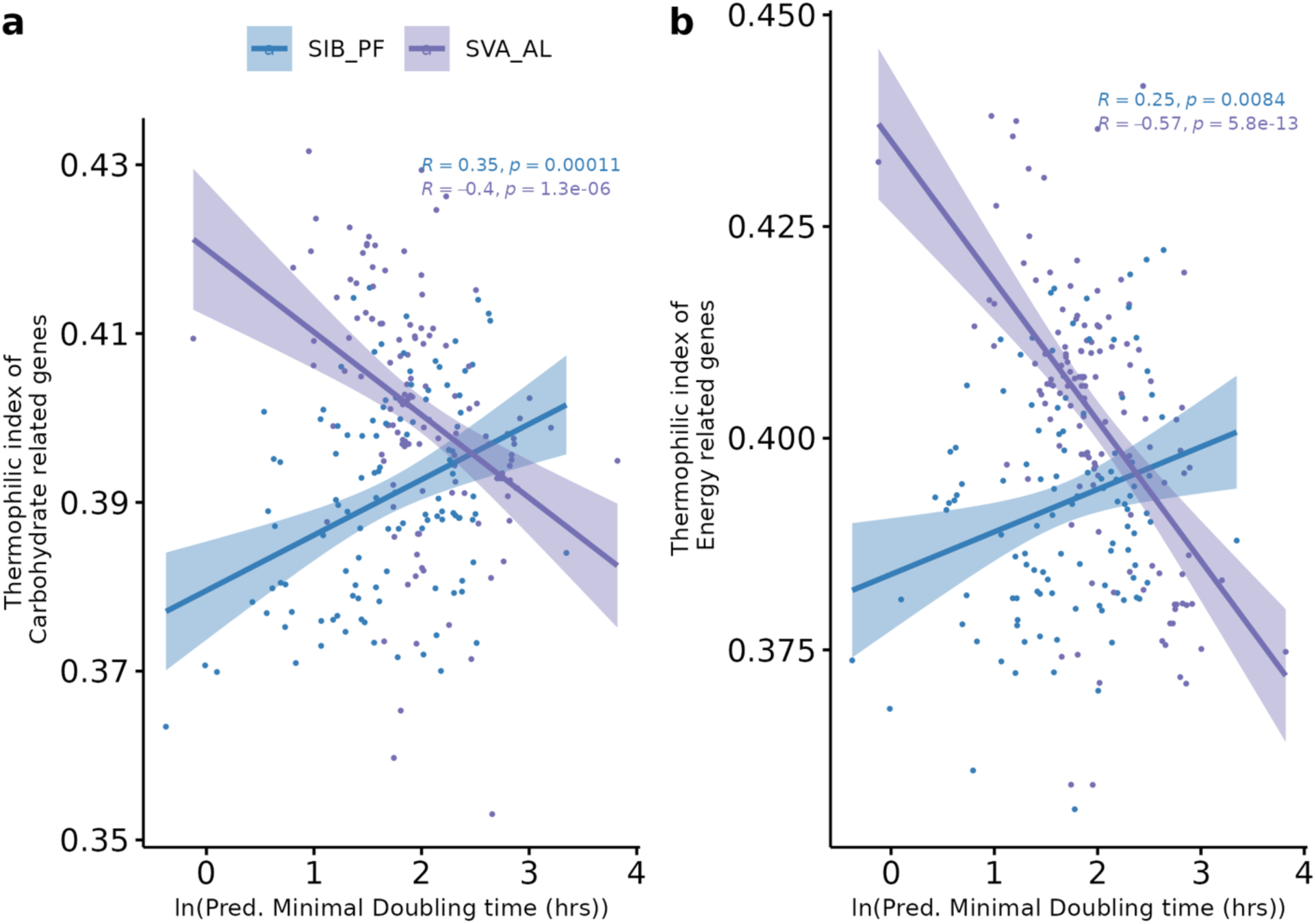
Correlations between predicted minimum doubling time and thermophilic index either specific to (a) annotated carbohydrate transport and metabolism genes (NCBI COG category ‘G’) or specific to (b) energy production and conversion genes (NCBI COG category ‘C’). The correlation analyses in a and b are two tailed. The shaded areas around the regression line show the 95% confidence interval.

### Thermophilic traits and minimum doubling time are independent of phylogeny in permafrost

To assess whether traits such as predicted optimal growth temperature and predicted minimum doubling time are influenced more by evolutionary lineage than environmental conditions, we measured the phylogenetic signal of predicted optimal growth temperature and predicted minimum doubling time using Pagel’s lambda (λ) (57). The phylogenetic signal evaluates whether a trait correlates with phylogenetic relatedness, indicating if shared evolutionary history affects its distribution. As shown in Table 1, λ values in permafrost were close to 0, indicating weak phylogenetic signal, while those in the active layer were close to 1, indicating a strong phylogenetic signal. This suggests that evolutionary lineage has a minimal effect on trait distribution in permafrost, potentially because traits shaped by environmental adaptation are under selective pressure that can override phylogenetic influence. In contrast, traits in the active layer show stronger phylogenetic influence, reflecting closer alignment with ancestral lineage.

**Table 1:** The phylogenetic signals of traits in permafrost (SIB_PF) and active layer (SVA_AL). PMDT indicated predicted minimum doubling time; POGT indicates predicted optimal growth temperature.

| SIB_PF |  |  |  | SVA_AL |  |  |
| --- | --- | --- | --- | --- | --- | --- |
| Traits | n | Pagel's $\lambda$ | p | n | Pagel's $\lambda$ | p |
| PMDT | 26 | < 0.001 | >0.05 | 27 | 0.999 | $1.337 \times 10^{-8}$ |
| POGT | 26 | < 0.001 | >0.05 | 27 | 0.999 | $3.114 \times 10^{-9}$ |

To further explore this, we applied phylogenetic generalized least squares (PGLS) regression (58) under a Brownian motion model to examine relationships between predicted optimal growth temperature and predicted minimum doubling time (Table 2). The slope from this model represents the phylogenetic rate of change in predicted optimal growth temperature as predicted minimum doubling time varies. In permafrost, a non-significant positive correlation was observed between predicted minimum doubling time and predicted optimal growth temperature, consistent with the weak phylogenetic signal. Conversely, in the active layer, predicted minimum doubling time showed a significant negative correlation with predicted optimal growth temperature, suggesting stronger phylogenetic influence on this trait. With nearly balanced sample sizes for permafrost (n = 26) and active layer (n = 27), our results suggest that traits in permafrost are likely shaped more by environmental adaptation rather than ancestral lineage, in contrast to traits in the active layer.

**Table 2:** PGLS regression of PMDT and POGT in permafrost (SIB_PF) and active layer (SVA_AL)

| Traits | SIB_PF |  |  |  |  | SVA_AL |  |  |  |  |
| --- | --- | --- | --- | --- | --- | --- | --- | --- | --- | --- |
|  | Slope | p | PBH | R<sup>2</sup> | Adj. R<sup>2</sup> | Slope | p | PBH | R<sup>2</sup> | Adj.R<sup>2</sup> |
| PMDT vs POGT | 2.99 | 0.112 | 0.223 | 0.128 | 0.082 | -5.19 | $4.27 \times 10^{-5}$ | $8.55 \times 10^{-5}$ | 0.495 | 0.475 |

### Thermophilic slow growers share few horizontally transferred genes with mesophilic permafrost microbes

Horizontal gene transfer (HGT) plays a significant role in the sharing of adaptive traits, particularly in permafrost environments where microbes have longer generation times (59). To investigate whether thermophilic slow growers engage in genetic interactions with other microbes in the permafrost, we categorized MAGs into four groups based on predicted optimal growth temperature and predicted minimum doubling time data: thermophilic fast growers, thermophilic slow growers, mesophilic fast growers, and mesophilic slow growers. For these purposes, “thermophilic” indicates a predicted optimum growth temperature above 40°C and “slow-grower” indicates a predicted minimum doubling time longer than five hours. In Siberian permafrost, 52 MAGs were identified as mesophilic fast growers, 50 MAGs as mesophilic slow growers, and 12 MAGs as thermophilic slow growers. In the Svalbard active layer, 23 MAGs were classified as mesophilic fast growers, 106 MAGs as mesophilic slow growers, and 4 MAGs as thermophilic fast growers. HGT predictions were made using MetaCHIP (56), which employs both phylogenetic and best-match approaches.

In permafrost communities, 703 genes were involved in HGT transfers, whereas in the active layer, only 63 genes were identified as HGT genes. In permafrost, the majority of HGT transfers occurred between mesophilic slow and fast growers, with only 7 HGT genes transferred within thermophilic slow growers. In contrast, within the active layer, thermophilic fast growers were more involved in HGT events, with 22 out of 63 HGT genes transferred within this group (Fig. 4). The fewer HGT events observed within the permafrost thermophilic group suggest that thermophilic traits may have evolved independently within this group.

**Fig. 4.**
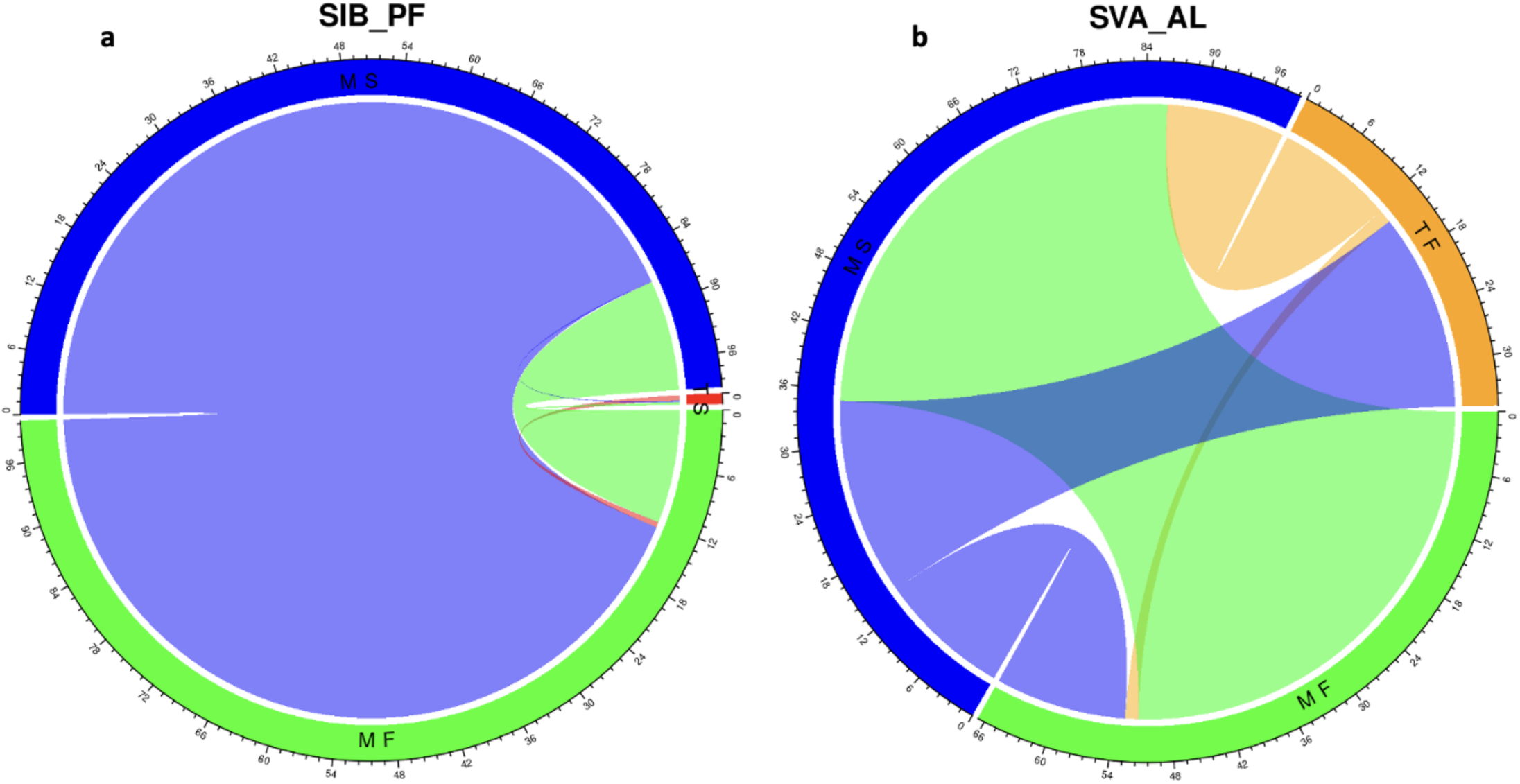
Predicted horizontal gene transfer events within the permafrost and active layer microbial communities. Bands connect the donors and recipients. The band’s color corresponds to the donors and the band width represents the number of HGTs. The customized groups include Thermophilic slow growers, TS (predicted optimal growth temperature > 40°C, predicted minimum doubling time > 5 hrs), Thermophilic fast growers, TF (predicted optimal growth temperature > 40°C, predicted minimum doubling time < 5 hrs), Mesophilic slow growers, MS (predicted optimal growth temperature between 20°C and 40°C, predicted minimum doubling time > 5 hrs), Mesophilic fast growers, MF (predicted optimal growth temperature between 20°C and 40°C, predicted minimum doubling time < 5 hrs). The 5 hrs cut off is based on gRodon classification of slow and fast growers [21]. The number outside the grid represents the matrix percentage of each donor.

## Discussion

### Permafrost shows thermophilic intrinsic adaptation to slow growth

Permafrost soils represent low-energy environments. Even though organic matter may be plentiful, cold temperatures and low water availability limit the rate at which metabolic reactions can proceed (60, 61). Growth rate predictions from gRodon suggest that the microbes that dominate both permafrost and overlying active layer soils grow slowly compared to microbes in a high-energy environment like the human gut (Fig. S1). While this is perhaps unsurprising, there has been disagreement in the literature about whether active-layer soil microbes are predominantly fast- or slow-growing (62, 63). Slow growth in permafrost is consistent with the low rates of metabolic processes, which reflects adaptation to the low energy environment (18, 27, 64–66).

It is likely that, for the taxa that are predicted to be slow growers in this analysis, predicted minimum doubling time is an underestimate of the true minimum doubling time, which itself is an underestimate of the *in situ* doubling time. gRodon uses codon usage bias to predict growth rate. Codon usage bias is mainly used by microbes to optimize translational efficiency, particularly in highly expressed genes (67–70). Thus, gRodon underestimates the minimum doubling times of taxa with minimum doubling times above ∼5 hrs (21). Despite gRodon’s tendency to underestimate growth rates of slow growers, it remains superior to alternative tools (49).

Previous work has shown that the true minimum doubling times of environmental microorganisms (which might be measured in pure cultures growing exponentially) are in turn substantially shorter than the *in situ* doubling times (71), since *in situ* doubling times are a function of a host of environmental factors, in addition to an organism’s biology. Thus, genomically predicted optimal growth rates may be a better indication of an organism’s preferred growth strategy. Despite the similar abundances of slow-growers between permafrost and active layers, the slow-growers in permafrost differed from those in the active layer in their apparent adaptation to temperature. In the active layer, in which episodic thawing and rainfall events cause microbes to episodically experience warmer temperatures and receive pulses of organic matter (24, 72, 73), faster-growing microbes were predicted via both TOME and the thermophilic index to be adapted to higher temperatures. This is consistent with a split between r-strategists, which capitalize on conditions in which fast growth is possible (warm temperature, substrate availability, water availability) and k-strategists, which persist effectively when fast growth rates are not possible, for instance between episodic warm or wet events (74).

The pattern in the permafrost was the opposite: taxa predicted to be slow growers were predicted to be adapted to higher temperatures. Furthermore, many microbes in permafrost had predicted optimal growth temperatures well above temperatures that they would ever experience *in situ*. It is common for microbes to have predicted optimal temperatures on the order of ∼10°C above typical *in situ* environmental temperatures (75–77), but many microbes in permafrost had optimal temperatures above 30°C, whereas *in situ* temperatures by definition never exceed ∼0°C (30).

The trends observed with the microbial thermophilic index and that of the carbohydrate and energy related genes were similar (Fig. 1c & Fig. 2). Also, the predicted optimal temperature for NDPK, the temperature optimum of which has previously been shown to reflect the environmental temperatures experienced by the host organism (78), match the same trend (Fig. 2d). What would be the value of adaptation to high temperatures in microbes that will never experience those temperatures?

We propose that the apparent high temperature optimum of slow-growing permafrost microbes is not an adaptation to high temperature *per se*, but rather an adaptation to extreme slow growth rates, the class of organism for which the term aeonophile has recently been proposed (22). Theoretical work indicates that as microbial metabolic rates approach zero, the energetic cost of protein repair increases as a fraction of a cell’s total energy requirements (79). All else being equal, a cell that turns its biomass carbon over more slowly must either have a lower fraction of active enzymes relative to inactive enzymes, or must have more stable enzymes, than a cell that turns its carbon over faster. The soil carbon turnover time in the permafrost is generally longer than that of the active layer across all ecosystems (80).

Adaptations for efficient energy acquisition and substrate utilization reflect strong selective pressures in permafrost, driving microbial community structure and function (81). Preliminary evidence from deep marine sediments, in which microbial metabolisms are also remarkably slow, suggests that secreted hydrolases are in fact unusually thermostable (82).

We note that we are agnostic as to whether the predicted thermophiles identified in these data sets are true thermophiles (or thermotolerant microbes), in the sense of being capable of growth or persistence at high temperatures. Rather, we argue that these microbes’ genomes code for the same traits that TOME (19) identifies as thermophilic, which primarily reflect protein stability. We suggest that enhanced protein stability also facilitates an ultra-slow-growth lifestyle at cold temperatures.

### Weak phylogenetic signal in traits correlation associated with permafrost

Our results indicate that the predicted optimal growth temperature and predicted minimum doubling time in permafrost communities exhibit weak phylogenetic signals. A weak phylogenetic signal indicates that closely related species may not share similar traits, implying that evolutionary history alone cannot account for the observed trait patterns. In such cases, ecological factors, particularly environmental adaptations, likely play a more critical role in shaping community composition (83, 84). Phylogenetic signal strength can vary across traits and scales. For instance, traits related to methane oxidation kinetics in methane-oxidizing bacteria display weak phylogenetic signals, highlighting the dominant influence of ecological adaptations (85). Similarly, weak phylogenetic signals observed in microbial traits related to resource use strategies suggest that environmental factors impact certain traits more than others (86–88).

This weak phylogenetic signal in permafrost microbes may also be linked to a high rate of horizontal gene transfer. Our analysis indicates that horizontal gene transfer occurs more frequently in permafrost than in the active layer (Fig. 4). Horizontal gene transfer can facilitate the spread of adaptive traits, especially in extreme environments like permafrost, where microbes have prolonged generation times (59). Moreover, horizontal gene transfer events in nature are linked to environmental dynamics and can contribute to the co-existence of microbial populations which further increase the microbial complexity (89, 90). However, in thermophilic slow growers within permafrost, we observed fewer genes that had been horizontally transferred both to and from other groups, suggesting an isolated evolutionary pathway (Fig. 4). It is unclear whether these thermophilic microbes are indigenous to permafrost or originated from thermophilic environments. Scarce hot springs and fumaroles within discrete permafrost-affected regions can support microbial communities, suggesting possible migration or gene flow (8, 91).

In permafrost or other cold environments, adaptation to slow growth may allow thermophilic microbes to be viable for hundreds to thousands of years (92, 93). This suggests at low temperatures, thermophilic traits may include nutrient utilization strategies that help counteract cold stress and low energy availability. Furthermore, slow growers may have acquired thermophilic traits to increase their resistance to the cold environments. The unique combination of low temperature and metabolic flexibility in permafrost may thus have shaped a microbial community that integrates traits for both thermophily and stability, reflecting the complex interplay of ecological and evolutionary processes in this extreme environment.

## Acknowledgments

The authors acknowledge funding through the Genomic Science Program of the U.S. Department of Energy, Office of Science, Office of Biological and Environmental Research (DE-SC0020369 to A.D.S. and T.A.V.) and from the National Science Foundation (OCE-2145434)

## Contributions

I.O. performed bioinformatic analyses and wrote the manuscript. T.A.V. and A.D.S. secured funding. All authors contributed to conceptualization of the research, writing, and editing the manuscript.

## Data availability

All metagenomic data sets are available in table S1 and as described by their original publications

